# Engineering genetically-encoded synthetic biomarkers for breath-based cancer detection

**DOI:** 10.1101/2021.09.01.456741

**Authors:** Ophir Vermesh, Aloma L. D’Souza, Israt S. Alam, Mirwais Wardak, Theresa McLaughlin, Fadi El Rami, Ataya Sathirachinda, John C. Bell, Michelle L. James, Sharon S. Hori, Eric R. Gross, Sanjiv Sam Gambhir

**Affiliations:** Department of Radiology, School of Medicine, Stanford University, Stanford, CA 94305, USA; Molecular Imaging Program at Stanford, Stanford University, Stanford, CA 94305, USA; Stanford Cardiovascular Institute, Stanford University, Stanford, CA 94305, USA; Stanford University Mass Spectrometry, Stanford University, Stanford, CA 94305, USA; Department of Microbiology and Immunology, Stanford University, Stanford, CA 94305, USA; Department of Biochemistry, Microbiology, and Immunology, University of Ottawa, Ottawa, Ontario, Canada; Department of Neurology and Neurological Sciences, School of Medicine, Stanford University, Stanford, CA 94305, USA; The Canary Center at Stanford, School of Medicine, Stanford University, Palo Alto, CA 94304, USA; Department of Anesthesiology, Perioperative and Pain Medicine, School of Medicine, Stanford University, Stanford, CA 94305, USA; Department of Bioengineering, Stanford University, Stanford, CA 94305, USA; Stanford Bio-X, Stanford University, Stanford, CA 94305, USA

**Keywords:** Cancer, breath, synthetic biomarker, metabolic engineering, early detection

## Abstract

Breath analysis holds great promise for rapid, noninvasive early cancer detection; however, clinical implementation is impeded by limited signal from nascent tumors and high background expression by non-malignant tissues. To address this issue, we developed a novel breath-based reporter system for early cancer detection using D-limonene, a volatile organic compound (VOC) from citrus fruit that is not produced in humans, in order to minimize background signal and maximize sensitivity and specificity for cancer detection. We metabolically engineered HeLa human cervical cancer cells to express limonene at levels detectable by mass spectrometry by introducing a single plant gene encoding limonene synthase. To improve limonene production and detection sensitivity twofold, we genetically co-expressed a modified form of a key enzyme in the cholesterol biosynthesis pathway. In a HeLa xenograft tumor mouse model, limonene is a sensitive and specific volatile reporter of tumor presence and growth, permitting detection of tumors as small as 5 mm. Moreover, tumor detection in mice improves proportionally with breath sampling time. By continuously collecting VOCs for 10 hours, we improve sensitivity for cancer detection 100-fold over static headspace sampling methods. Whole-body physiologically-based pharmacokinetic (PBPK) modeling and simulation of tumor-derived limonene predicts detection of tumors as small as 7 mm in humans, equivalent to the detection limit of clinical imaging modalities, such as PET, yet far more economical.

**Significance Statement:** We developed a breath-based reporter system using the plant terpene, D-limonene – a volatile secondary metabolite that gives citrus fruit its characteristic scent but is not produced in human tissues – as a biomarker for early cancer detection. Results from this study could pave the way for *in vivo* gene delivery and tumor-specific expression of exogenous volatile cancer reporters with broad applicability to the early diagnosis of a wide variety of cancers.

## Introduction

Breath analysis provides rapid and non-invasive biomolecule detection, with great promise for early cancer detection and surveillance^1,2^. The human body emits hundreds of volatile organic compounds (VOCs) – organic molecules that readily vaporize at room temperature – in the breath^3^. Breath, a less complex matrix than blood and other bodily fluids, can be sampled easily, painlessly, and inexpensively^1^. Moreover, breath can be directly analyzed using real-time mass spectrometry, reducing the need for sample storage and processing^1^. While no single VOC can reliably signal cancer presence on its own, VOC signatures or “breathprints” have been reported that can distinguish a number of cancers – including lung, colon, breast, and prostate cancers – from benign disease and healthy controls in relatively small study populations^4–6^. However, as with liquid biopsies^7–14^, clinical implementation of breath VOCs for early cancer detection is limited by low or variable signal from cancer cells and high background signal from nonmalignant tissues^3,5^. Furthermore, identification of cancer-specific VOC signatures has been impeded by statistical overfitting – a common pitfall in early stage ‘omics approaches due to typically small study populations relative to the numerous endogenous parameters analyzed – limiting their generalizability.^15^

Engineered synthetic reporters provide an innovative solution to overcome the detection limitations of endogenous biomarkers.^16–21^ By effecting diseased cells to express an exogenous biomarker that is not naturally produced in human tissues, background signal from non-diseased tissues is minimized, thereby maximizing sensitivity and specificity. Moreover, exogenous reporters from biochemical classes that are orthogonal to the human metabolome can easily be distinguished from the complex milieu of endogenous molecules by mass spectrometry. Furthermore, detection of a single exogenous biomarker that uniquely signals disease presence avoids the statistical challenges associated with endogenous VOC analysis. Recent synthetic strategies include exogenous protein biomarkers encoded on *in vivo*-delivered DNA vectors and selectively secreted into the blood by cancer cells, as well as nanoparticles that release a volatile compound in the breath to signal lung infection or inflammation^17,21^. Genetically-encoded synthetic biomarkers have practical and theoretical advantages, including: 1) integration with clinically established *in vivo* gene delivery methods, including those used in vaccine development (viral vectors, liposomes, and minicircles)^22–24^; 2) selective expression in many cancer types using tumor-specific promoters and tumoritropic or tumor-targeted vectors^17,25,26^; and 3) continuous expression throughout the lifetime of the cancer, which can enable repeat monitoring after a single administration. However, there have been no reports thus far of strategies that genetically encode synthetic biomarkers for breath-based detection of cancer.

Here, we combine the high specificity and sensitivity of an exogenous cancer biomarker with the speed, simplicity, and non-invasive nature of breath VOC detection. To genetically encode a VOC biomarker in cancer cells that is distinct from endogenous VOCs, we looked to plants. Humans and plants share a common cholesterol biosynthesis pathway, but in plants this pathway also generates terpenes, the volatile compounds that attract pollinators and protect from herbivorous insects and pathogens^27–29^. We hypothesized that the mammalian cell’s cholesterol biosynthetic machinery could be exploited to produce plant volatiles by genetically introducing the appropriate exogenous enzymes (**Fig. 1**). While many plant volatiles require multiple biosynthetic steps, only a single enzyme, limonene synthase (LS)^30^, bridges the cholesterol biosynthesis pathway with production of limonene^27^, the monoterpene that gives citrus fruits their characteristic scent. Limonene is already used clinically (for example, to treat gallstones and heartburn), has chemopreventive and chemotherapeutic effects in many types of cancers, and is safe at oral doses as high as 100 mg/kg (~7 g for an average 70 kg adult)^31,32^. Due to its wide industrial use, metabolic engineering approaches for increasing limonene biosynthesis have been extensively studied in microbial systems and plants^29,33–36^, and have the potential to be adapted to human cancer cells for breath-based diagnosis and eventually – at high expression levels – for therapy. We demonstrate that limonene can be genetically expressed in human cancer cells and can report on early tumor presence and growth in a xenograft mouse model. We also extrapolate our VOC-based detection to humans using a whole-body physiologically-based pharmacokinetic (PBPK) model of VOC biodistribution, metabolism, and exhalation.

**Fig. 1.**
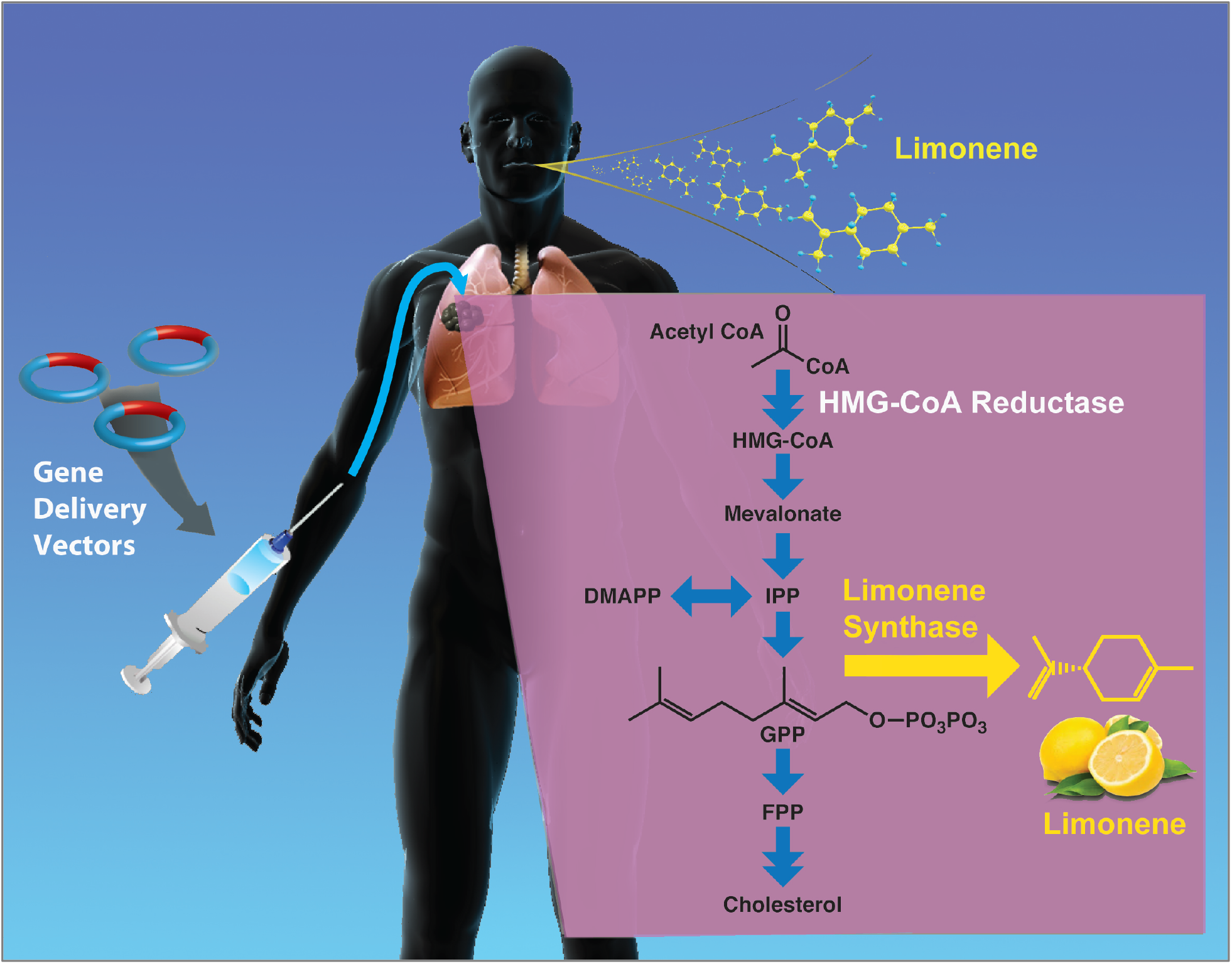
Envisioned cancer reporter strategy using a plant-based volatile organic compound. A cancer patient undergoing surveillance or a healthy subject undergoing cancer screening is administered a gene delivery vector (minicircle, liposome, or adenovirus) encoding an exogenous plant enzyme – driven by a pan-tumor-specific promoter – which catalyzes production of a plant volatile organic compound (VOC) specifically in cancer cells that is not otherwise produced endogenously. The VOC diffuses into the bloodstream and is transported to the lungs, where it is exhaled in the breath and detected by a breath analyzer (mass spectrometer or electronic nose sensor array), uniquely signaling the presence of cancer and overall tumor burden. While a lung tumor is shown above to illustrate the concept, we expect this strategy to be generalizable to many cancer types. **Inset:** Expressing a plant VOC in a human cell. Plants and humans share a conserved metabolic pathway for cholesterol production (blue arrows) but in plants, terpene synthases divert part of this metabolic stream towards production of volatile organic compounds that attract pollinators and protect from herbivorous insects, parasites, and pathogens. Selective expression of terpene synthases, such as limonene synthase (yellow arrow), in human cancer cells could enable these cells to produce plant VOCs that are detectable in breath, serving as highly specific cancer reporters. Substrates in the cholesterol biosynthetic pathway: HMG-CoA, 3-hydroxy-3-methylglutaryl coenzyme A; DMAPP, dimethylallyl pyrophosphate; IPP, isopentenyl diphosphate; GPP, geranyl diphosphate; FPP, farnesyl pyrophosphate.

## Results

### Limonene expression and detection in cultured tumor cells

HeLa cells were transfected with a vector containing LS and eGFP genes under the control of a single CAG promoter (**Figs. 2A, 2B**). Antibiotic selection and FACS sorting for high eGFP expressers yielded a stable cell line containing limonene synthase (HeLa-LS) (**Fig. 2C**). To maximize limonene production in cultured HeLa-LS cells, we targeted a key regulatory enzyme of the mevalonate pathway, HMG-CoA reductase (HMGR)^37^. Truncation of HMGR by deletion of its N-terminal regulatory domain renders it insensitive to feedback inhibition by downstream metabolites, augmenting flux through the mevalonate pathway and increasing the availability of limonene precursors. Previous studies in bacteria and yeast engineered to produce limonene have shown that expression of truncated HMGR (tHMGR) can markedly increase limonene production^33,34^. We transfected HeLa-LS cells with a plasmid encoding human tHMGR and turbo red fluorescent protein (tRFP) under the control of an EF1α promoter (**Figs. 2A, 2B**). Antibiotic selection and FACS sorting for high expression of tRFP yielded a stable cell line expressing both eGFP and tRFP (**Fig. 2C**), and containing both LS and tHMGR (HeLa-LS-tHMGR). Solid phase microextraction (SPME) fibers^5,38^ were used to sample the culture headspace (i.e. the air above the cells) in flasks containing confluent stably transfected cells (**Fig. 2A**). Gas chromatography-mass spectrometry (GC-MS) analysis of the fibers showed a mass spectrum closely matching the limonene standard, with both exhibiting the characteristic ion peaks for limonene (m/z = 68, 93, and 136)^39^ at the same relative ratios (**Fig. 2D**) and identical chromatogram retention times (**Fig. 2E**).

**Fig. 2.**
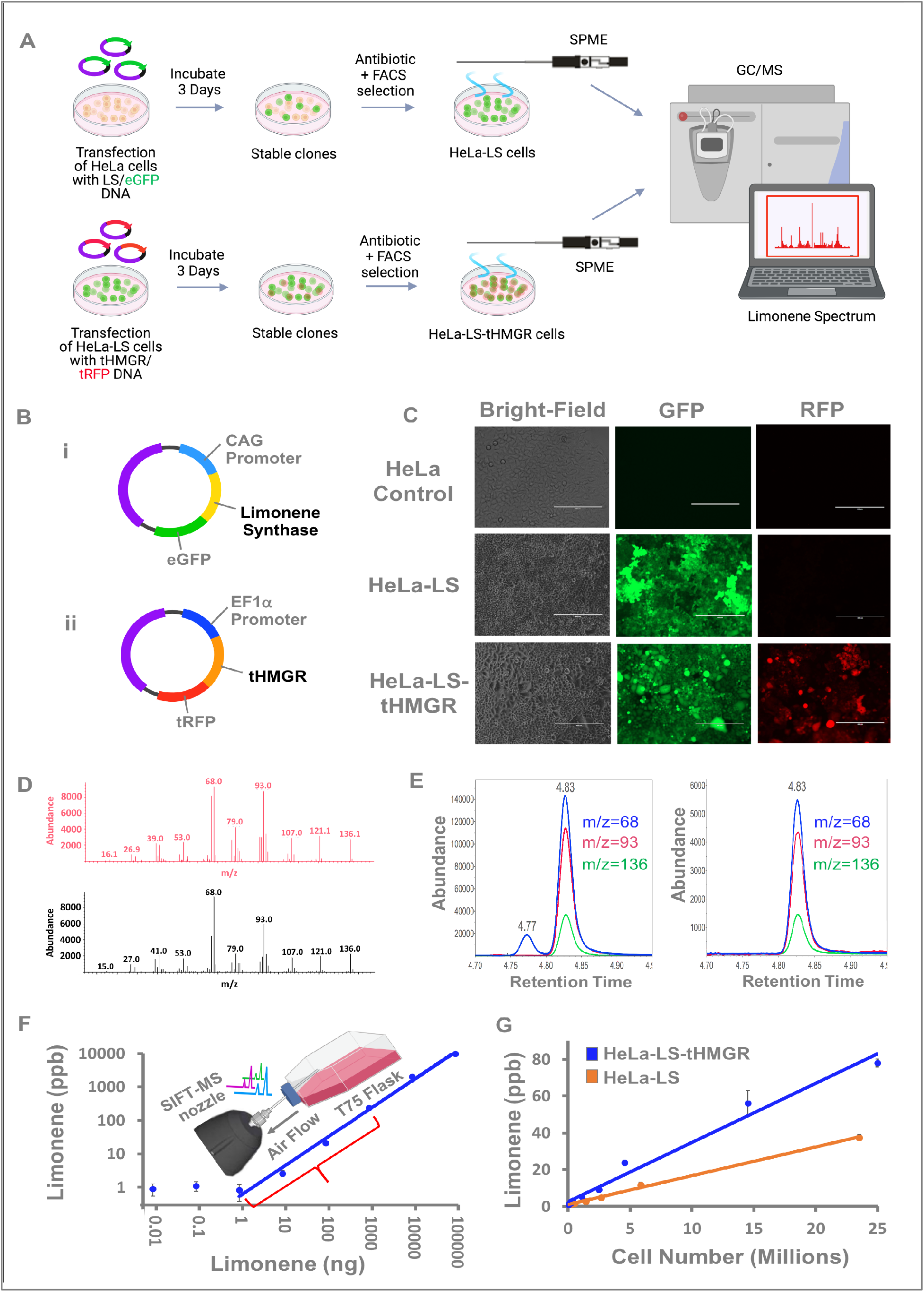
Vector design, transfection, and limonene production by HeLa cells. (**A)** Schematic of experimental methodology. **(Top)** Cultured HeLa cells were transfected with a vector containing LS and eGFP genes under the control of a CAG promoter. Antibiotic and FACS selection for stably transfected clones (sorting on eGFP-expressing cells) resulted in a HeLa cell line containing both LS and eGFP (HeLa-LS-eGFP cells, subsequently referred to as HeLa-LS cells). (**Bottom**) HeLa-LS cells were subsequently transfected with a vector containing the tHMGR and tRFP genes under the control of an EF1α promoter. Antibiotic and FACS selection (based on dual expression of eGFP and tRFP) resulted in a HeLa cell line containing LS, tHMGR, eGFP, and tRFP (HeLa-LS-tHMGR-eGFP-tRFP, subsequently referred to as HeLa-LS-tHMGR). Solid phase microextraction (SPME) fibers were used to sample the culture headspace of confluent stably transfected HeLa-LS and HeLa-LS-tHMGR cells for 30 minutes, and were then analyzed for limonene by GC-MS. **(B) (i)** Piggybac transposon DNA vector containing truncated limonene synthase (LS) and enhanced green fluorescent protein (eGFP) driven by a CAG promoter, and puromycin resistance gene driven by a CMV promoter. **(ii)** Piggybac transposon DNA vector containing truncated HMG CoA reductase (tHMGR) and turbo red fluorescent protein (tRFP) driven by an EF1α promoter, and hygromycin resistance gene driven by a CMV promoter. **(C)** Bright-field and fluorescence images showing HeLa-LS and HeLa-LS-tHMGR cells after antibiotic selection and FACS sorting, compared with untransfected control HeLa cells. Scale bar = 200 um for HeLa control and 400 μm for HeLa-LS and HeLa-LS-tHMGR. **(D)** Mass spectrum from an SPME fiber exposed to the headspace of confluent HeLa-LS cells **(top, red)** compared with the reference spectrum of limonene from a mass spectrum library (Mnova database) **(bottom, black).** Note the characteristic peaks at m/z = 68, 93, and 136. **(E)** Selected ion monitoring (SIM) mode chromatogram of an SPME headspace sample from HeLa-LS cells (left) and from a pure limonene standard (right), showing matching ion ratios and retention times. **(F)** Calibration curve relating headspace limonene concentration as measured by SIFT-MS to the quantity of limonene spiked into culture media in a T75 flask (*y* = 0.62*x*^0.86^, R^2^=0.99). Over the range of limonene production by cultured cells (1 to 1000 ng, red bracket), the relationship is well-modeled by *y* = 0.28*x* (R^2^=0.99). **(G)** Headspace concentration of limonene as a function of cell number for HeLa-LS (*y* = [1.56×10^−6^]*x* + 1.06, R^2^ = 0.99) and HeLa-LS-tHMGR cells (*y* = [3.21×10^−6^]*x* + 2.70, R^2^ = 0.98) after incubation at 37°C for 24 hours. Limonene measured from HeLa-LS-tHMGR cells was approximately double that from HeLa-LS cells over the cell density range examined.

### Quantification of limonene from transfected cells

We further confirmed the presence of headspace limonene using selected ion flow tube mass spectrometry (SIFT-MS), which affords continuous, real-time VOC detection with quantification down to the parts-per-billion leve^13,5,40^. To obtain quantitative measurements of headspace limonene, we created a calibration curve for limonene (10 pg to 100 μg) spiked into media within a 280 mL T75 flask (**Fig. 2F**). Headspace concentrations increased as a function of *x*^0.86^ for limonene quantities within the range of 1 ng to 100 μg (R^2^ = 0.99) and demonstrated a nearly linear dependence with limonene quantities ranging from 1 ng to 1 μg (R^2^ = 0.99). The limit of detection (LOD) for limonene by SIFT-MS was 1.8 ng, corresponding to 0.5 ppb in the headspace. Next, we sought to quantify limonene generated by transfected HeLa cells over a 24-hour period. Limonene production increases linearly over a range of 45,000 to 25 million cells for both HeLa-LS (R^2^=0.99) and HeLa-LS-tHMGR (R^2^=0.99), with LODs of 360,000 cells and 107,000 cells, respectively, as compared to undetectable limonene levels in untransfected HeLa cells (**Fig. 2G, Supplementary Calculations**). For the largest number of HeLa-LS cells tested, a confluent culture of 23.5 million cells, the headspace limonene concentration was 38±2 ppb, corresponding to 131 ng of limonene or an average of ~5.6 fg per cell per day. For the largest number of HeLa-LS-tHMGR cells tested, 25 million cells, the headspace limonene concentration was 78 ± 2 ppb, corresponding to 277 ng of limonene or an average of ~11 fg per cell per day. The slope of the best-fit line for HeLa-LS-tHMGR cells was twice that for HeLa-LS cells (3.2×10^−6^ vs. 1.6×10^−6^), demonstrating that HeLa-LS-tHMGR cells generated double the amount of limonene as HeLa-LS cells.

### Quantification of limonene emitted from limonene-injected and tumor-bearing mice

Having observed robust limonene expression in transfected HeLa cells in culture, we then tested the feasibility of detecting limonene in exhaled breath from rodents. We created a standard curve relating limonene concentration in chamber headspace to the quantity of limonene spiked into 0.5-L chambers. To determine the fraction of limonene in mice that is emitted into the headspace, we injected mice intraperitoneally with different quantities of a limonene standard solution (from 0.01 μg to 1 mg) and placed individual mice in a closed chamber for 15 minutes, at which point headspace limonene concentrations were measured by SIFT-MS (**Fig. 3A, 3B**). Using the standard curve, we determined the mass of limonene exhaled by mice at each quantity injected, and calculated the fraction exhaled. At the LOD (0.5 ppb), limonene in the chamber headspace became detectable when 2.3 ng had been spiked into the chamber, whereas limonene evolving from mice only became detectable at an injected dose of 450 ng (**Fig. 3B, Supplementary Calculations**). A comparison of the graphs for these two conditions shows that only ~0.5% of limonene at each injected dose was emitted into the chamber headspace within 15 minutes of injection. We therefore expected mice bearing limonene-producing tumors to emit a similar fraction into the chamber headspace over this time period. Assuming the limonene production rate in cell culture to be an upper bound on the expected cellular limonene production rate in tumor-bearing mice, we calculated that large tumors with diameters of at least 3.4 cm (4 billion cells) would be required in order to reach the detection limit of SIFT-MS within 15 minutes (**Supplementary Calculations**). To test this, we implanted one million HeLa-LS or HeLa-LS-tHMGR cells subcutaneously into each flank of immunocompromised nude mice and monitored them using SIFT-MS at 5 weeks post-implantation. Consistent with our calculations, we found that no limonene was detected in the chamber headspace even when up to 4 mice with a combined tumor burden of ~4 cm^3^ were contained in a single chamber.

**Fig. 3.**
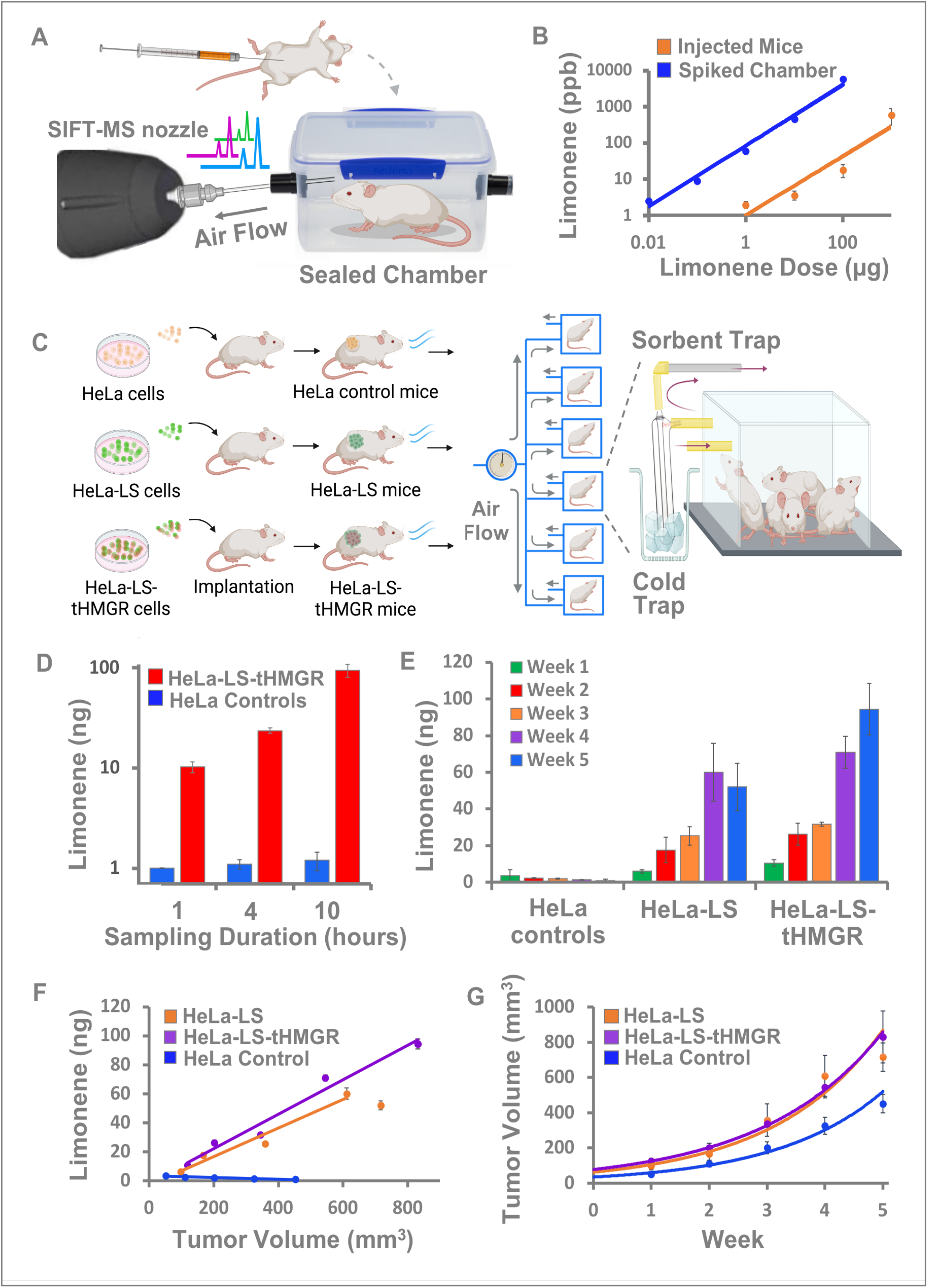
Limonene detection from mice. **(A)** Intraperitoneal injection of limonene into a mouse, placement of the mouse in a sealed 0.5-L chamber, and SIFT-MS analysis of chamber air after 15 minutes. **(B)** Limonene concentration in chamber headspace as a function of limonene dose injected intraperitoneally into mice (*y* = 1.01*x*^0.82^, R^2^ = 0.89) or spiked directly into a chamber containing 10 ml of water (*y* = 83.83*x*^0.84^, R^2^ = 0.99). Only ~0.5% of limonene injected into mice was detected in chamber air at 15 minutes. Each data point represents mean ± SD for n = 3 mice (one mouse per chamber). **(C)** Ten-week-old athymic nude mice were inoculated subcutaneously in both flanks with either HeLa-LS, HeLa-LS-tHMGR, or untransfected control HeLa cells. Tumor progression in the 3 groups was followed over a 5-week period with weekly measurements of tumor size and collection of mouse VOCs using a specially-designed mouse chamber setup in which highly purified air was continuously flowed into 6 one-liter mouse chambers (4 mice per chamber) in parallel at 100 mL/min. Air exiting the chamber was flowed through a cold trap to eliminate moisture and then through a sorbent trap containing Tenax resin to capture VOCs from the mice. The sorbent traps were subsequently analyzed by GC-MS. **(D)** Limonene signal in HeLa-LS-tHMGR mice increases with sampling time, whereas limonene signal in control mice remains below the detection limit (<2.3 ng), demonstrating that signal-to-noise ratio and sensitivity can be increased by increasing the sampling time. **(E)** Five-week follow-up study of grouped mice implanted with HeLa-LS, HeLa-LS-tHMGR, and untransfected control HeLa cells. Limonene production increased with time post-implantation for HeLa-LS and HeLa-LS-tHMGR mice and was detectable above background at one week post-implantation in HeLa-LS-tHMGR mice. Peak limonene production in HeLa-LS-tHMGR mice was significantly greater that in HeLa-LS mice (94 ± 14 ng vs. 60 ± 16 ng, p = 0.049). **(F)** Limonene production by HeLa-LS and HeLa-LS-tHMGR mice increases approximately linearly with tumor volume over the first 4 weeks of the study. HeLa-LS: *y* = 0.10*x* - 3.2, R^2^ = 0.95. HeLa-LS-tHMGR: *y* = 0.12*x* – 1.76, R^2^ = 0.97. Limonene was undetectable in control mice with untransfected HeLa tumors. **(G)** Tumor growth rates for all three groups were modeled assuming monoexponential growth. HeLa-LS-tHMGR: *y* = 77.3e^0.48*t*^, R^2^ = 0.99. HeLa-LS: *y* = 62.2e^0.53*t*^, R^2^ = 0.96. HeLa controls: *y* = 34.5e^0.54*t*^, R^2^ = 0.98. Each bar or data point for limonene quantity represents mean ± SD for 3 chambers of 4 mice each (n = 12 mice). “Tumor volume” refers to the average tumor volume in a single mouse (i.e. group tumor volume divided by 4).

To increase sensitivity for detecting limonene from tumor-bearing mice, we built a specially-designed experimental setup in which highly purified air is continuously flowed through a mouse chamber and exits through an air sampling tube containing a sorbent material (Tenax TA) that traps VOCs, thereby pre-concentrating them for subsequent GC-MS analysis. Compared to SPME fibers, sorbent traps contain much larger quantities of sorbent material and therefore have higher extraction capacities^5^. Six one-liter chambers were set up in parallel to allow for multiple simultaneous experiments (**Figs. 3C, S1**). We placed groups of HeLa-LS-tHMGR mice and control mice bearing untransfected HeLa tumors at 5 weeks post-implantation into side-by-side chambers, with 4 mice per chamber (average tumor volume per mouse: 1.2 ± 0.2 cm^3^), and sampled the chamber headspace (100 mL/min airflow) for 1, 4, or 10 hours. In the experimental group, limonene was detectable in chamber air at all sampling durations. Increasing the sampling duration from 1 hour to 4 hours enabled 2.3-fold greater limonene collection (10 ng to 23 ng), and an increase to 10 hours enabled 9.4-fold greater limonene collection (10 ng to 94 ng) (**Fig. 3D**). Limonene levels for control mice were below 1 ng at all sampling durations. Therefore, we showed that increased signal-to-background was achievable simply by sampling the chamber headspace for a longer time. By integrating limonene signal over a number of hours, the sorbent trap method improves detection sensitivity 100-fold compared to direct SIFT-MS measurements in sealed unventilated chambers (**Supplementary Calculations**), where measurements are limited to only a few minutes before mice become hypoxic. To maximize our sensitivity, we chose 10-hour sampling times for all subsequent mouse experiments.

We next sought to determine the minimum tumor size at which limonene is detectable and to evaluate whether tumor growth could be monitored via exhaled limonene alone. HeLa-LS, HeLa-LS-tHMGR, and control mice (bearing untransfected HeLa tumors) were monitored over a 5-week period. Groups of four mice per chamber (n=3 chambers per cohort) were tested once a week for total limonene released into chamber air during a 10-hour period. At week one post-implantation of tumor cells, total evolved limonene from the HeLa-LS-tHMGR cohort (11 ± 2 ng) was statistically higher compared to the HeLa-LS (6 ± 1 ng, p = 0.049) and control mouse groups (4 ± 3 ng, p = 0.025) (**Figs. 3E, S2**). At this time, the average tumor volume per mouse was 0.12 cm^3^, 0.10 cm^3^, and 0.05 cm^3^, for HeLa-LS-tHMGR, HeLa-LS, and control mice, respectively (**Figs. 3F, G**). Average limonene per mouse in the HeLa-LS-tHMGR group (~2.7 ng) at week one was very close to our calculated detection limit (2.3 ng), which suggests that the minimum detectable tumor size by VOC sampling is close to 0.1 cm^3^, or 4.6-mm diameter (corresponding to approximately 10 million HeLa cells, see **Supplementary Calculations**). Evolved limonene from HeLa-LS mice was not statistically different from controls (p = 0.26) at week one. Thus, the expression of tHMGR by limonene-producing cancer cells aided in detecting tumors earlier relative to mice with limonene-producing tumors that did not express tHMGR, as expected based on the higher production of limonene by HeLa-LS-tHMGR cells in culture. By the second week, evolved limonene was statistically higher in both HeLa-LS-tHMGR (26.3 ± 6.0 ng, p = 0.025) and HeLa-LS mice (17.6 ± 6.9 ng, p = 0.025) than in control mice (2.3 ± 0.3 ng) (**Figs. 3E, S2**), at an average tumor volume per mouse of 0.2 cm^3^, 0.18 cm^3^, and 0.1 cm^3^, respectively (**Figs. 3F, G**).

Limonene emitted from HeLa-LS and HeLa-LS-tHMGR mice increased linearly with tumor volume over 4 and 5 weeks post-implantation, respectively (**Fig. 3F**). Limonene evolution was higher in HeLa-LS-tHMGR mice than in HeLa-LS mice throughout the study, though this difference was statistically significant only in weeks 1 and 5. Limonene evolution from HeLa-LS and HeLa-LS-tHMGR mice peaked in weeks 4 and 5 at 60 ± 16 ng and 94 ± 14 ng, respectively (when tumor burden per mouse was 0.6 ± 0.1 cm^3^ and 0.8 ± 0.2 cm^3^, respectively). This plateau in HeLa-LS mice corresponded with a leveling off in tumor growth (i.e. no statistical change) from weeks 4 to 5 (**Fig. 3F**). At week 5, mice were humanely euthanized due to tumor size. Tumor growth rate, *k*, was slightly greater in control mice (*k* = 0.54) than in HeLa-LS-tHMGR (*k* = 0.48, p = 0.049), whereas it was not statistically different between HeLa-LS-tHMGR and HeLa-LS mice (*k* = 0.53, p = 0.13) or between HeLa-LS and control mice (p = 0.51) (**Fig. 3G**). Limonene quantities collected from HeLa control mice at each time point were very similar to blank chambers without mice, with a range of <1 ng to 4 ng (**Fig. S3**). These values likely represent ambient limonene that was degassing from the chamber walls, given that limonene levels both from control mice and blank chambers were below the detection limit by the end of the 5-week study. Moreover, limonene was not detected above background in chambers containing only mouse diet gel or bedding. Therefore, we concluded that the only sources of limonene in HeLa-LS-tHMGR and HeLa-LS mice were the tumors. The average percentage of tumor limonene exhaled in the breath over all weeks was calculated at 5.2% ± 1.5% and 7.6% ± 3.1% for HeLa-LS-tHMGR and HeLa-LS mice, respectively (**Supplementary Calculations**, **Tables S3-S7**).

### Whole-body physiologically-based pharmacokinetic (PBPK) model for VOC detection

To predict the smallest tumor size that would be detectable in humans using a sorbent trap, we developed and simulated a whole-body physiologically-based pharmacokinetic (PBPK) model for limonene disposition in humans (**Fig. 4A, Tables S1-S2**). Our multi-compartment model shows that in humans, 1.3% of tumor-derived limonene would be emitted in the breath (**Fig. 4B**). This result agrees with prior literature showing that, due to rapid biodistribution (particularly to adipose tissue) and metabolism, ~1% of absorbed limonene is eliminated unchanged in exhaled breath^31,32,41,42^. Given the LOD of the sorbent trap method (2.3 ng), we calculated that a human tumor would need to produce at least 177 ng (1.3% of 177 ng is 2.3 ng) of limonene over a 10-hour period to be detectable in the breath. Assuming a limonene production rate similar to HeLa-LS-tHMGR cells, tumors would become detectable when they reached 7 mm in size (**Supplementary Calculations**). Substituting limonene with a VOC that has low fat-to-blood (K_f:b_) or blood-to-air (K_b:a_) partition coefficients could further improve detection sensitivity. For example, our PBPK model shows that if instead of limonene (K_f:b_ = 140, K_b:a_ = 42)^2^, a VOC reporter with partition coefficients like those of 2-butanone (K_f:b_ = 0.75, K_b:a_ = 215)^43^ was chosen, 30% of it would be exhaled in the breath (**Fig. 4C**); this could permit detection of tumors that are ~30-fold smaller by volume (2.3 mm diameter) at the same VOC expression level. Our PBPK modeling approach can also be extended to other potential VOC candidates for cancer detection.

**Fig. 4.**
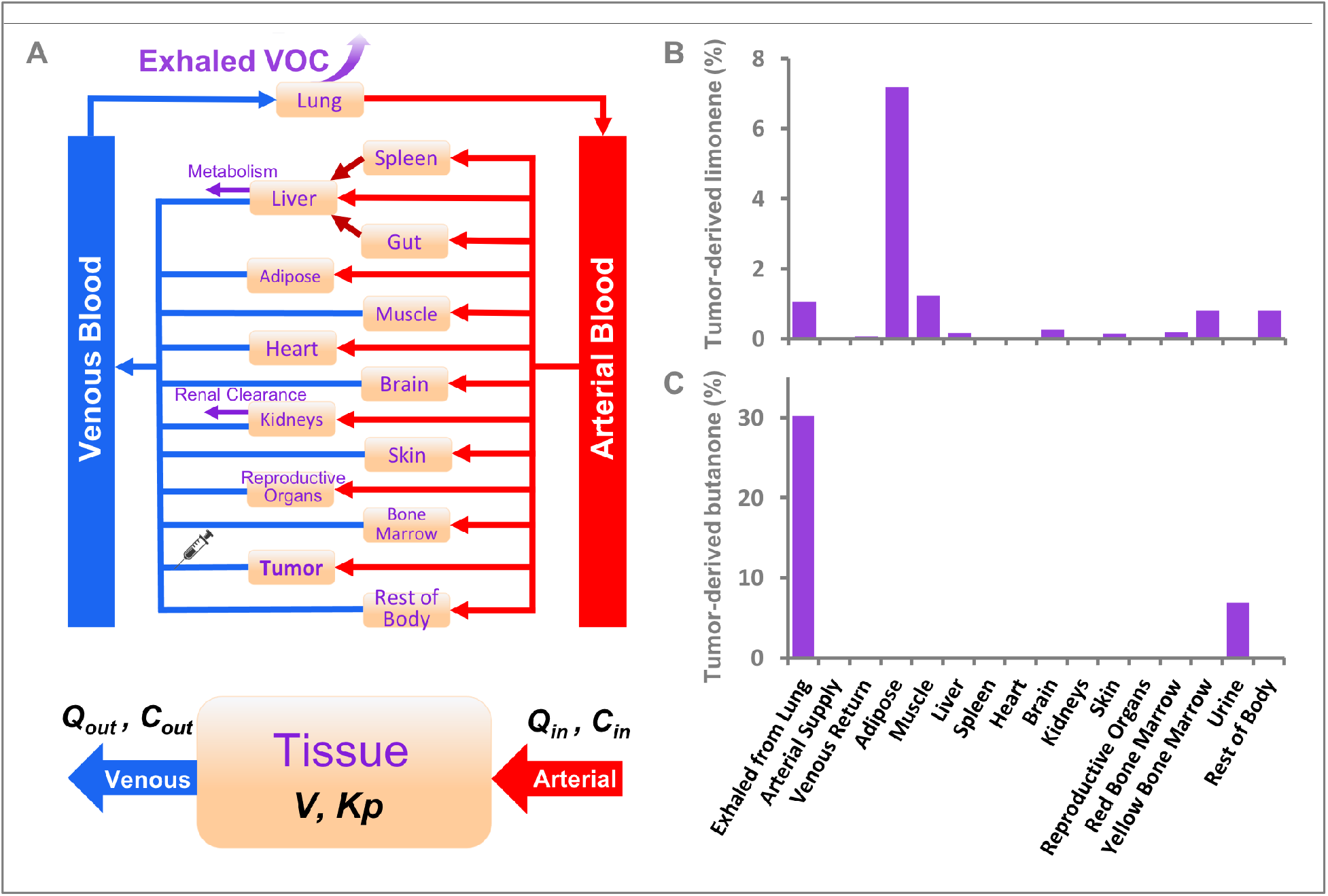
Human whole-body PBPK model for synthetic VOC biomarkers. **(A)** Schematic of the human whole-body PBPK model for two synthetic VOC biomarkers (limonene and butanone), indicating the major tissues and organs modeled and key input parameters. The tissues are linked by the arterial and venous blood compartments, and each tissue is characterized by an associated blood flow rate, volume, tissue-to-plasma partition coefficient, and permeability. ***Q_in_*** and ***Q_out_*** represent blood flow rate into and out of the tissue, respectively; ***C_in_*** and ***C_out_*** represent the VOC concentration of blood flowing into and out of the tissue, respectively; ***V***, tissue volume; ***K_p_***, tissue-to-plasma partition coefficient of VOC. Tumor release of the VOC (limonene or butanone) into the blood was simulated as a constant intravenous infusion. Limonene **(B)** and butanone **(C)** distribution in humans at steady state assuming a constant intravenous infusion by tumors producing these synthetic VOC biomarkers. Note that the percentage of limonene is highest in adipose tissue (7%) with only a small fraction (1.3%) exhaled in the breath. The majority of limonene (>85%) is metabolized. The percentage of butanone is highest in exhaled breath (30%), followed by urine (7.5%), with negligible quantities in the tissues. The remainder (>60%) is metabolized.

## Discussion

Here, we report a novel strategy for sensitive and specific breath-based cancer detection that uses limonene, a plant terpene, as an exogenous VOC reporter. First, we demonstrated stable heterologous expression of limonene, as validated by mass spectrometry, in a cultured HeLa human cervical cancer cell line transfected with a plasmid encoding the plant enzyme limonene synthase. We also demonstrated that genetically co-expressing a modified key mevalonate pathway enzyme, tHMGR, can double limonene expression in HeLa cells, thereby improving detection sensitivity for these cells in culture and *in vivo*. We then validated limonene as a sensitive and specific volatile reporter of tumor presence and growth in a xenograft mouse model after subcutaneous implantation of limonene-expressing HeLa cells, and showed that limonene can be detected when tumors are as small as 120 mm^3^ (~5 mm diameter). Using human whole-body PBPK modeling, we predicted that tumor-derived limonene would be detectable in human breath from a tumor as small as 7 mm in diameter.

The attributes of an ideal breath-based cancer reporter include: 1) safety; 2) low background (little or no endogenous production in the body); 3) specificity (high expression in cancer cells relative to nonmalignant cells; 4) good deliverability from tumor to the lungs (high air-to-plasma and plasma-to-tissue partition coefficients, and low fat-to-plasma partition coefficient); 5) little or no metabolism to non-volatile compounds; 6) abundance of biochemical precursors; and 7) few enzymatic steps (facilitating ease of design and efficient production in mammalian cells). Identification of a VOC reporter that meets all of these criteria is challenging. Limonene is an attractive candidate because it is safe, not endogenously produced in human tissues, and requires only a single enzymatic step for expression in mammalian cells. For any given VOC, factors such as deliverability to the lungs and rate of metabolism are dependent on the VOC’s structure and intrinsic molecular properties and cannot easily be controlled, except by switching to a different VOC reporter altogether. However, various strategies can be implemented to increase detection sensitivity, mitigate other sources of background, increase the availability of biochemical precursors, and achieve tumor-specific expression, as we discuss below.

A key finding from our work was that sensitivity for detecting small tumors could be dramatically improved via sorbent trapping of limonene from a mouse chamber over extended periods of time. A 10-fold increase in sampling time from one to 10 hours yielded a concomitant increase in limonene signal from HeLa/LS mice, while background limonene levels for HeLa control mice remained below the detection limit. Further integrating limonene signal over even longer periods of time may enable detection of even smaller tumors. The breath can provide an essentially limitless sample volume for this purpose since the total volume sampled can be increased simply by breathing into a sorbent trap for a longer period of time. By contrast, there are practical limits to the amount of blood that can be drawn from a patient in a given time (10% of total blood volume per month)^44^.

While extending sampling time increases limonene signal, sensitivity can also be improved by minimizing background noise. Due to widespread industrial use and production of the S-enantiomer by many plant species, limonene is present at 0.1 to 2 ppb in ambient air^31,32^. In this study, background limonene levels in control and empty chambers decreased each week as the chambers were cumulatively exposed to highly pure air over a longer period of time, further removing residual limonene from the chamber walls. Thus, further improvements in LOD may be possible by pre-purging chambers with air for longer periods of time (e.g., a few weeks) or running experiments in glass chambers, which do not readily adsorb hydrophobic VOCs. In the clinical scenario, human subjects would be placed in a room with highly pure air or would breathe through a one-way filter cartridge to prevent contamination of inhaled air by ambient limonene. Exhaled air would pass through an exhaust valve directly into a sorbent tube, which would subsequently be analyzed offline by GC-MS. The small filter cartridge/sorbent tube assembly could be worn portably to passively collect limonene over a few hours as the subject goes about their day or at night while sleeping. Subjects would need to avoid wearing perfumes or consuming citrus prior to undergoing testing. The presence of limonene in the breath at screening or surveillance would then prompt clinical imaging studies, such as PET or MRI, in an attempt to spatially localize the tumor. Monitoring of VOC reporter levels could also be used to assess response to therapy inexpensively and more frequently than is practical or economical with *in vivo* imaging in patients with metastatic disease or large disease burden.

For cancer screening and early detection, we envision targeting expression of the limonene synthase gene to cancer cells using novel and clinically relevant gene delivery approaches including minicircles^17,24^. Smaller tumors could then be detected by employing strategies that increase cellular reporter gene production. Engineering 10- or 100-fold increases in tumor limonene production could permit detection of human tumors as small as 3 mm (below the detection limit of clinical PET imaging) or 1.5 mm, respectively (**Supplementary Calculations**). High cancer reporter expression can obviate time-consuming pre-concentration steps by achieving VOC levels in the breath that are directly detectable by rapid breath analysis platforms such as SIFT-MS, an electronic nose, or eventually a portable breath analyzer for routine at-home cancer monitoring. Clinically, certain aromas in breath have long been recognized by unaided olfaction as telltale signs of various pathologic states, including the ketone odor in diabetic ketoacidosis and the ammonia odor in chronic kidney disease^3^. Very high expression levels (>10 ppb) could likewise allow limonene to be detectable in the breath by the human nose (i.e. cancer diagnosis by a quick “sniff test” at the doctor’s office).

Metabolic engineering strategies that have been used for large-scale microbial biosynthesis of limonene and other natural products^27,33–36,45,46^ can potentially be adapted to scale up tumor cell production of VOC reporters in human cells in culture and *in vivo*. The production of limonene is mainly limited by the availability of its direct biochemical precursor, geranyl diphosphate (GPP), in the cytosol^29,34^. In mammalian cells, GPP is a relatively short-lived metabolic intermediate because the synthase enzyme that converts isopentenyl diphosphate (IPP) to GPP then rapidly converts GPP to farnesyl pyrophosphate (FPP). Truncated forms of GPP- and FPP-synthases have been investigated in engineered microbes that predominantly produce GPP, increasing monoterpene production^33–35,47^. Another strategy that has successfully increased terpene production in microbes is to introduce additional copies of mevalonate pathway genes into cells. Moreover, since the methylerythritol 4-phosphate (MEP) pathway provides most of the carbon flux for terpene production in plants, heterologous expression of these enzymes (without the PSP) could potentially increase limonene production in human cells as well^35,47–49^. Combinations of genes could in principle be delivered on the same vector as part of a comprehensive *in vivo* gene delivery approach.

Future work will test strategies for *in vivo* gene delivery and expression of our VOC reporter. Lung cancer would intuitively be an ideal initial clinical target for sensitive breath-based detection because of the close proximity of lung tumors to exhaled air. Further, using our PBPK model, we plan to identify additional candidate VOC reporters with improved deliverability to the lungs. For a VOC reporter that is not consumed in the diet or present environmentally, its background levels would be limited only by the tumor-specificity of the gene delivery approach or gene expression method. A potential hurdle to clinical implementation of any exogenously-derived reporter is the development of host immunity to the foreign enzyme, which could render subsequent delivery of the gene ineffective. However, techniques for de-immunization, in which epitope sequences are modified to minimize immunogenicity without altering protein structure or function, have long been used in biologic drug design and can similarly be applied to limonene synthase^50^. Tolerogenic sequences can also be introduced to promote immune tolerance to exogenous enzymes^51,52^. Collectively, we anticipate that these efforts will help advance this technology to facilitate cancer screening and early detection in the clinic.

## Materials and Methods

### Study approval

All procedures performed on animals were approved by Stanford University’s Institutional Animal Care and Use Committee (Stanford, CA, USA) and in compliance with the guidelines for humane care of laboratory animals.

### Vector design (HeLa-LS and HeLa-LS-tHMGR)

The sequence for R-limonene synthase was codon-optimized for expression in human cells using the GenSmart Codon Optimization tool (GenSript, Pascataway, NJ). The plastid signaling peptide (PSP), which functions independently of enzyme activity to localize R-limonene synthase to plastids in plants, was excluded as it impairs proper folding in other expression systems^33,53^. The truncated limonene synthase (LS) gene exhibited markedly higher limonene production in bacterial culture compared to the full-length gene^34^, and was therefore used for the duration of the study. Mammalian PiggyBac transposon gene expression vectors coding for LS or a modified hydroxy-3-methylglutaryl-CoA reductase (tHMGR) were designed using VectorBuilder (https://en.vectorbuilder.com/design.html) and constructed by Cyagen Biosciences. The vector encoding LS also contained the gene for enhanced green fluorescent protein (eGFP) linked by a P2A ribosomal skip sequence^54^, with both genes driven by the same CAG promoter. This vector also contained a puromycin resistance gene driven by a CMV promoter for antibiotic selection. The vector encoding tHMGR also contained the gene for turbo red fluorescent protein (tRFP) linked by a P2A ribosomal skip sequence, with both genes driven by the same EF1α promoter. This vector also contained a hygromycin resistance gene driven by a CMV promoter for antibiotic selection.

### Cell culture

HeLa cells (American Type Culture Collection, Manassas, VA) were cultured in Dulbecco’s Modified Eagle Medium (DMEM) media supplemented with penicillin-streptomycin and 10% fetal bovine serum (FBS) (ThermoFisher, Waltham, MA). Cells were verified to be free of mycoplasma contamination using the MycoAlert Mycoplasma Detection Kit (Lonza, Allendale, NJ) and passaged when reaching 80% confluence.

### HeLa cell transfection

HeLa cells were transfected with a LS-encoding vector using Lipofectamine 2000 (Invitrogen, Carlsbad, CA). The ratio of the LS vector to a helper plasmid containing the transposase gene was 1:1 (0.8 μg of each per well in a 12-well plate) in Gibco Opti-MEM Reduced Serum media (ThermoFisher, Waltham, MA). Stable transfection was assessed qualitatively under fluorescence microscopy by the visual presence of high GFP expression in cells at days 3-4 post-transfection. Cells subsequently underwent antibiotic selection and multiple rounds of fluorescence-activated cell sorting (FACS) to select for high-expressing GFP subclones and were tested for limonene production as described below. This cell line was named HeLa-LS. Transfection of limonene-producing cells with a tHMGR-encoding vector (HeLa-LS-tHMGR) was accomplished in a similar manner, with hygromycin B (ThermoFisher, Waltham, MA) used for antibiotic selection of stable cells, and with FACS selection performed by gating on RFP (**Fig. 2A, B**).

### Fluorescence-activated cell sorting

Roughly 1-2 million confluent stably transfected cells were sorted on a FACS Aria II or Influx sorter (Becton Dickinson, San Jose, CA). The gating strategy included forward scatter (FSC) and side scatter (SSC) gating, doublets and dead cell exclusion, and selection for the top 1-2% highest expressers of eGFP for LS-expressing cells, or tRFP for pre-sorted LS-expressing cells transfected with the vector containing the tHMGR gene.

### Cell culture headspace sampling (SPME)

Stably transfected HeLa-LS or HeLa-LS-tHMGR cells were grown to confluence in T75 flasks (MIDSCI, St. Louis, MO) at 37°C. The 24-gauge needle of a solid-phase microextraction (SPME) assembly (Sigma Aldrich, St. Louis, MO) was inserted through the screw cap septum of the T75 flask and the 65-μm PDMS/DVB fiber was deployed for 30 minutes to sample the cell culture headspace. The fiber was withdrawn and adsorbed VOCs were analyzed by gas chromatography/mass spectrometry (GC/MS).

### Gas Chromatography-Mass Spectrometry

Analysis of SPME fibers was performed on an Agilent 7890/5975 GC/MS instrument (Agilent Technologies, Santa Clara, CA) at the Stanford Mass Spectrometry Facility. One microliter of sample was injected through an SPME inlet guide (Supelco, Bellefonte, PA) into the GC injection port, equipped with a Thermogreen LB-2 pre-drilled septum (Supelco) and deactivated glass inlet liner (Supelco), and run in pulsed splitless mode. Helium was used as the carrier gas with a constant flow rate of 1.6 mL/min and velocity of 27.8 cm/s through an Agilent DB-WAX column (60 m x 250 μm x 0.25 μm). The initial oven temperature was held at 40°C for 2 minutes, increased at a rate of 2°C/min up to 72°C, then ramped at 40°C/min to 220°C. Total run time was 21.7 minutes. Initial scans were run in full scan mode at m/z 10-400. Subsequently, samples were run in selected ion monitoring (SIM) mode, targeting the characteristic ion peaks for limonene: m/z 68, 93, and 136.

### Quantitation of limonene production in HeLa cells

Prior to cell studies, a calibration curve was generated. Serial dilutions of pure limonene (Sigma Aldrich, St. Louis, MO) in ethanol were prepared in Eppendorf tubes and spiked into 10 mL of media (DMEM with 10% FBS) to final concentrations ranging from 0.01 ng to 100 μg in T75 flasks with screwcap septa (MIDSCI, St. Louis, MO). The flasks were manually agitated for 10 seconds and the screw cap septum was punctured by a needle. The flask headspace was sampled for 20 seconds at least 3 times per concentration using selected ion flow mass spectrometry (SIFT-MS, Syft Technologies, Christchurch, New Zealand) with a helium gas carrier. Limonene detection was performed by soft-ionization using H_3_O^+^ (m/z, 137; branching ratio, 68%; reaction rate, 2.6×10^−9^ cm^3^/s), NO^+^ (m/z, 136; branching ratio, 88%; reaction rate, 2.2×10^−9^ cm^3^/s) and O2+ (m/z, 93; branching ratio, 29%; reaction rate, 2.2×10^−9^ cm^3^/s) to calculate limonene concentration in real-time. After establishing the calibration curve, HeLa-LS and HeLa-LS-tHMGR cells were spiked into 10 mL media (DMEM with 10% FBS) in varying numbers ranging from 20,000 to 10 million cells in T75 flasks. The flasks were incubated at 37°C for 24 hours, after which headspace limonene concentrations were measured using SIFT-MS. The cells were then harvested and counted with cell numbers at harvest ranging from ~45,000 to 25 million.

### Quantitation of limonene evolution from limonene-injected mice

Prior to mouse studies, a calibration curve was generated. Known limonene quantities (10 pg to 100 μg) were added to 10 mL of water in 0.5-mL chambers (Kent Scientific, Torrington, CT). The chambers were capped, briefly agitated, and allowed to sit for 15 minutes to equilibrate. The chamber inlet was then uncapped and the headspace was sampled by SIFT-MS for limonene. After establishing the calibration curve, serial tenfold dilutions of limonene in ethanol were prepared and a twenty-microliter volume of each solution (1 to 1000 μg limonene) was injected intraperitoneally into immunocompromised nude mice. The injection site was rinsed thoroughly under warm water for 15 seconds to remove possible limonene residue from the skin. Each mouse was then placed in a closed 0.5-L chamber for 15 minutes, at which point the chamber inlet was uncapped and the headspace was sampled by SIFT-MS for 20 seconds.

### Xenograft tumor mouse model

Ten-week-old athymic nude (nu/nu) mice (Charles River Laboratories, Wilmington, MA) were inoculated subcutaneously in both flanks with either HeLa-LS, HeLa-LS-tHMGR, or untransfected control HeLa cells (1 million cells in 100 μL of Matrigel [ThermoFisher, Waltham, MA] into each flank). Prior to each experiment, mouse tumors on both flanks were measured via caliper and the tumor length (L), width (W), and depth (D) were recorded to calculate tumor volumes (V) using the equation for volume of an ellipsoid: 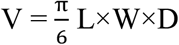.

### Mouse chamber/sorbent trap assembly

Six one-liter chambers (Braintree Scientific, Braintree, MA) were operated in parallel for simultaneous mouse limonene measurements (**Fig. S1**). The outlet of each chamber was connected in series via tygon tubing to a glass condenser (25 mL impinger, SKC Ltd., UK) on ice (cold trap) and then to a sorbent tube containing Tenax TA resin (Markes International Ltd., UK) that traps and concentrates VOCs. The cold trap prevents moisture from soaking the sorbent resin. The inlet of each chamber was connected in series to a sacrificial Tenax sorbent tube, which served to purify inflowing air, and an upstream ¼” stainless steel metering valve (Swagelok Company, Solon, OH) that individually controlled air flow into each chamber. The metering valves to all six chambers were connected via reducing unions, union tees, and 1/8” copper tubing to a benchtop pressure regulator (Markes International Ltd., UK, U-GAS03) set to 5 psi, which was connected via a single copper line to a compressed gas cylinder containing highly pure air (Vehicle Emission Grade Air, Airgas Inc., Radnor, PA) set to 20 psi. For ease of cleaning the induction chambers between experiments, the tygon connections to inlet and outlet components were interrupted by ¼” snap-on/snap-off fasteners (Thermoplastic Quick Couplings, Omega Engineering Inc, Norwalk, CT).

### Operation of chamber/sorbent trap assembly for VOC sampling from tumor mice

Prior to initial mouse experiments, the induction chambers were flushed with highly pure air at 100 mL/min for 3 days. On the evening prior to experiments, 40 mL of mouse bedding and diet gel (CearH2O, Portland, ME) were placed in each chamber, and air flow was continued overnight (~10 hours) with the Tenax tubes connected to measure the background limonene levels in empty chambers. On the day of experiments, mice were pre-hydrated with a subcutaneous injection of 0.5 mL sterile saline. Air flow was continued for 30 minutes after mice were placed in the induction chambers to remove any ambient limonene entering while the chambers were briefly open. Tenax tubes were then replaced. A flow meter (Ellutia 7000, Ellutia Ltd, UK) measured the air flow exiting each Tenax tube and the pin valves were tuned to achieve an air flow rate of 100 mL/min. When removing or replacing the screw caps on Tenax tubes, care was taken to keep the tube ends covered with a clean glove to prevent contamination from ambient air. Air was flowed continuously for the duration of the experiments (10 hours). After each experiment, mice were placed back in their cages. The chambers were then rinsed with water, 70% ethanol, and dried before highly pure air flow was resumed at 20 mL/min to maintain low background limonene levels in the chambers prior to subsequent experiments. Upon completion of mouse experiments, Tenax tubes were stored on ice and shipped to ALS Environmental (Simi Valley, CA) for thermal desorption and GC/MS analysis.

### Whole-body physiologically-based pharmacokinetic (PBPK) modeling and simulation of VOC disposition in humans

GastroPlus® version 9.8 (Simulations Plus, Inc, Lancaster, CA), embedded with the PBPKPlus™ module, was used to run the human whole-body PBPK simulations based upon a 30-year-old, 70 kg adult male. Multi-compartment modeling of organs relevant to the biodistribution, metabolism, and excretion of candidate VOCs was used in combination with a mechanistic multi-compartment pulmonary model to predict the fraction of tumor-derived VOC within exhaled breath. The pulmonary model describes the lungs as a collection of four compartments: an extra-thoracic compartment (nasopharynx, oropharynx, and larynx); a thoracic compartment (trachea and bronchi); a bronchiolar compartment (bronchioles and terminal bronchioles); and an alveolar-interstitial compartment (respiratory bronchioles, alveolar ducts and sacs, and interstitial connective tissue)^55,56^. The whole-body PBPK model was used to predict the pharmacokinetic profile and tissue distribution of a drug or compound as a function of time based on the compound’s physicochemical properties and disposition data. Physicochemical properties of compounds that were not available in the literature were predicted with the ADMET Predictor software tool version 8.5 (Simulations Plus, Lancaster, CA). Tumor release of limonene or butanone into the blood was simulated as a constant intravenous infusion. Simulations were run to predict the fraction of tumor-derived VOC biomarkers in exhaled breath and in the major tissues as a function of time.

### Statistical Analysis

Statistical analysis was performed using Microsoft Excel for Mac version 16.30 (Microsoft Corporation, Redmond, WA). Data presented represents the mean ± standard deviation (SD) unless otherwise noted. Best-fit lines for calibration data were obtained by linear regression. R^2^ correlation coefficients were calculated to assess goodness of fit. Limonene quantities from SIFT-MS measurements were interpolated from the best-fit curves. Limit of detection (LOD) and limit of quantification (LOQ) values were calculated using the following: 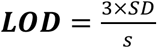 and 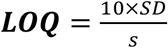, as per the International Conference on Harmonisation (ICH) and United States Pharmacopoeia (USP), where *SD* is the standard deviation of the residuals (vertical distance from a data point to the linear regression line) or of blank samples, and *s* is the slope of calibration curve at low concentrations^57^. For hypothesis testing, P values were calculated using a one-sided non-parametric test (Mann-Whitney U test). A P-value < 0.05 was considered statistically significant.

## Supporting information

VOC - Supplementary Information

## Acknowledgments

We thank Claire Keller, Wade Bontempo, David Wevill, and Paul Tobias at Markes International for important insights and onsite assistance with the design of the experimental setup for continuous air sampling from mice; Tim D. Knaak and Marty Bigos from the Stanford Shared FACS facility for their assistance with cell sorting; and Michael Bolger, John DiBella, Jim Mullin and Dr. Jessica Spires at Simulations Plus, Inc. for their expert advice and for lending us access to the GastroPlus software. GC/MS was performed using instruments at the Stanford University Mass Spectrometry Facility and at the Canary Center at Stanford. Figure graphics were created with BioRender.com.

## Funding

This research was supported by a grant from the Canary Foundation, the National Institutes of Health NIGMS GM119522 (ERG), and NIH S10 Shared Instrument Grant S10RR025518-01.

## Dedication

In loving memory of the late Dr. Sanjiv Sam Gambhir – professor, mentor, and friend – whose extraordinary impact and enduring vision live on in our hearts and minds and will continue to guide our future work for years to come.

## Notes

### Competing Interest Statement

The authors have declared no competing interest.

