## Supplementary material for "Engineering genetically-encoded synthetic biomarkers for breath-based cancer detection": VOC - Supplementary Information

**Affiliations:**

**Supplementary Calculations**

**Limit of Detection (LOD) for HeLa-LS-tHMGR cells:**

| # Cells Harvested | Predicted Limonene Concentration (ppb): Calibration Curve* | Actual Limonene Concentrations (ppb): Replicate 1 | Replicate 2 | Replicate 3 |
|---|---|---|---|---|
| 45967 | 0.22 | -0.01 | -0.09 | 0.25 |
| 88233 | 0.42 | 0.42 | 0.41 | 0.45 |
| 20300 | 0.97 | 0.67 | 1.00 | 0.89 |
| 378000 | 1.80 | 1.83 | 1.69 | 2.13 |

| Residuals (Actual minus Predicted): Replicate 1 | Replicate 2 | Replicate 3 |
|---|---|---|
| -0.23 | -0.31 | 0.03 |
| 0.00 | -0.01 | 0.03 |
| -0.29 | 0.03 | -0.08 |
| 0.03 | -0.12 | 0.33 |

- For LOD calculations, a best-fit curve through the four lowest-concentration data points yielded $y = (4.77\times10^{-6})x$ with $R^2 = 0.97$.
- Standard deviation of residuals = 0.17 ppb.
- Slope of calibration curve = $4.77 \times 10^{-6}$.
- $LOD = 3 \times SD/\text{slope} = 3 \times 0.17/(4.77 \times 10^{-6}) =$ **107,000 cells**.
- This corresponds to a concentration of $y = (4.77 \times 10^{-6}) \times 107{,}000 = 0.51$ ppb.
- Using the equation relating headspace concentration in a T75 flask to the amount spiked ($y = 0.28z$), 0.51 ppb corresponds to a total of 1.82 ng limonene produced.
- Combining the equations for: 1) headspace concentration as a function of cell number for the full range of cell numbers tested ($y = [3.2\times10^{-6}]x + 2.70$) with 2) headspace concentration as a function of spiked limonene quantity in ng ($y = 0.28z$), we obtain: $0.28z = (3.2\times10^{-6})x + 2.7$, so $z = (11.4\times10^{-6})x + 9.6$. The slope is the amount of limonene (in ng) produced per cell in a 24-hour period: $11.4\times10^{-6}$ ng, or ~11 fg/cell/ day.

**Limit of Detection (LOD) for HeLa-LS cells:**

| | Predicted Limonene Concentration (ppb) | Actual Limonene Concentrations (ppb) | | |
|---|---|---|---|---|
| # Cells Harvested | Calibration Curve* | Replicate 1 | Replicate 2 | Replicate 3 |
| 190500 | 0.27 | 0.36 | 0.08 | 0.16 |
| 503333 | 0.71 | 0.79 | 0.60 | 0.86 |
| 1425000 | 2.02 | 1.79 | 1.79 | 1.75 |
| 2651667 | 3.77 | 3.91 | 3.74 | 4.00 |

| Residuals (Actual minus Predicted) | | |
|---|---|---|
| Replicate 1 | Replicate 2 | Replicate 3 |
| 0.09 | -0.19 | -0.11 |
| 0.07 | -0.11 | 0.15 |
| -0.23 | -0.23 | -0.27 |
| 0.15 | -0.03 | 0.24 |

- For LOD calculations, a best-fit curve was drawn through the four lowest-concentration data points, yielding $y = 1.42 \times 10^{-6}x$ with $R^2 = 0.99$.

- Standard deviation of residuals = 0.17 ppb.

- Slope of calibration curve = $1.42 \times 10^{-6}$.

- $LOD = 3 \times SD/\text{slope} = 3 \times 0.17/(1.42 \times 10^{-6})$ = **360,000 cells**.

- This corresponds to a concentration of $y = (1.42 \times 10^{-6}) \times 359{,}155 = 0.51$ ppb.

- Using the equation relating headspace concentration in a T75 flask to the amount spiked ($y = 0.28z$), 0.51 ppb corresponds to limonene production of 1.82 ng.

- Combining the equations for: 1) headspace concentration as a function of cell number for the full range of cell numbers tested ($y = [1.56\times10^{-6}]x + 1.06$) with 2) headspace concentration as a function of spiked limonene quantity in ng ($y = 0.28z$), we obtain: $0.28z = (1.56\times10^{-6})x+1.06$, so $z = (5.6\times10^{-6})x+3.8$. The slope is the amount of limonene (in ng) produced per cell in a 24-hour period: $5.6\times10^{-6}$ ng, or ~5.6 fg/cell/ day.

- Based on these results, the limit of detection of the SIFT-MS instrument for limonene is 0.5 ppb above baseline, corresponding to 1.82 ng of limonene in a T75 flask.

- Similarly, a limonene LOD by SIFT-MS of 0.51 ppb corresponds to 2.3 ng of limonene in the empty 0.5-L chamber ($y = 83.83x^{0.84}$) and 0.43 µg of limonene injected into mice in 0.5-L chambers ($y = 1.01x^{0.82}$).

**Limit of Detection (LOD) of limonene from tumor mice in sorbent trap experiments**
For sorbent trap experiments, we calculated LOD using the equation:
$LOD = 3 \times SD_{blank}$ where $SD_{blank}$ is the standard deviation of a blank or control sample.
The standard deviation for untransfected HeLa control mice was 0.75.
Therefore, $LOD = 3 \times 0.75 = 2.3$ ng.

**Fraction of injected limonene exhaled in the breath**
To determine the fraction of limonene emitted into chamber air after injection into mice, we can rearrange the variables in the equations above so that limonene spiked into empty chambers $x_1$ or injected into mice $x_2$ is expressed in terms of headspace concentration, and solve for ($x_1/x_2$).

$y_1$=83.83 ($x_1$)^0.84 (empty chamber equation)
$y_2$=1.01 ($x_2$)^0.82 (i.p. injection in mouse)

For a given chamber headspace concentration $k=y_1=y_2=1$ ppb:
1 ppb = 83.83 ($x_1$)^0.84 → $x_1$=0.0051 µg
1 ppb = 1.01 ($x_2$)^0.82 → $x_2$=0.99 µg
So $x_1/x_2$ = 0.5%.

For $k=y_1=y_2$=10,000 ppb:
10,000 ppb = 83.83 ($x_1$)^0.84 → $x_1$=296.58 µg
10,000 ppb = 1.01 ($x_2$)^0.82 → $x_2$=74,606.87 µg
So $x_1/x_2$ = 0.4%.

**Comparison of sorbent trap and SIFT-MS sensitivity**
At week 5, HeLa-LS-tHMGR mice exhaled 94 ± 14 ng of limonene over a 10-hour period. This limonene quantity in the 0.5 L chambers used for the SIFT-MS experiments corresponds to a headspace concentration of 34 ppb at room temperature.

Concentration (ppb) = Concentration (ng/L) x [Molar volume (L)/Molecular Mass (g/mol)], where molar volume at 1 atm and room temperature is 24.45 L and limonene molecular weight is 136.24.

Concentration (ppb) = (94 ng/0.5L) x [24.45/136.24] = 33.7 ppb

Groups of 4 mice show sign of hypoxia in unventilated 0.5 L chambers after 6 minutes, at which point they have exhaled 0.94 ng of limonene, corresponding to a headspace concentration of 0.34 ppb, which is below the LOD of our SIFT-MS instrument (0.5 ppb). Therefore, the 100-fold improvement in limonene signal and sensitivity (34 ppb vs. 0.34 ppb) is due to the ability to collect sample over a 100-fold longer duration with the sorbent trap method.

**Fraction of tumor limonene exhaled in the breath**

The fraction of limonene produced by tumors that is exhaled in the mice was estimated by dividing the actual amount of limonene detected from tumor-bearing mice in the chambers (using Tenax tubes) by the predicted limonene production for a given tumor volume. The limonene production rates in cultured tumor cells (5.6 fg/cell/day for HeLa-LS cells and 11.1 fg/cell/day for HeLa-LS-tHMGR) were assumed to be upper bounds on the rate of limonene production by tumor cells *in vivo* since culture conditions are optimized with regard to nutrients and gas exchange. An estimate of an upper bound on the number of cells within a given tumor volume can be obtained by assuming that the entire tumor volume is occupied by tumor cells. The cell number can be calculated by dividing the total volume of the tumor by the average volume of individual tumor cells. While the number of cells per volume of tumor is frequently estimated to be $10^9$ cells/$cm^3$ [6,7], this assumes a 10 µm diameter cell. The average diameter of HeLa cells is approximately 20 µm, corresponding to ~$10^8$ cells per $cm^3$ volume[49]. Therefore, the fraction of tumor limonene exhaled in the breath by HeLa-LS-tHMGR mice in week 1 can be calculated as follows:

Total number of tumor cells on average in groups of four HeLa-LS-tHMGR mice in week 1 is tumor volume x $10^8$ cells/$cm^3$ = 0.492 $cm^3$ × $10^8$ cells/$cm^3$ = 49.2 million cells. Assuming an upper bound of 11.1 fg/cell/day (or 4.6 fg/cell in 10 hours), the maximum amount of limonene produced by this number of cells in a 10 hour experiment is 49.2 million x 4.6 fg = 227.7 ng. The actual amount of limonene exhaled in 10 hours from these mice was 7.1 ng. Therefore, at least 3.1% of the limonene produced by tumors was exhaled in the breath (**Tables S3-S7**).

**Predicted smallest detectable tumor in humans using limonene as a VOC reporter.** Given that only 1.3% of limonene produced by tumors *in vivo* would be exhaled in the breath, and given that the LOD of the sorbent trap method is 2.3 ng (for over 60 L of air sampled), a human tumor would need to produce 177 ng of limonene over the measurement duration to be detectable in the breath. At a measurement duration of ten hours as in our mouse experiments and assuming limonene production rates of 11.1 fg/cell/day as in HeLa-LS-tHMGR cells, we can calculate the minimum number of tumor cells and minimum tumor size detectable as follows:

$$177 \times 10^{-9} \text{ grams}/[11.1 \times 10^{-15} \text{ grams/cell} \times (10 \text{ hours}/24 \text{ hours})] = 38.3 \text{ million cells}$$

Given that 1 $cm^3$ of HeLa-LS-tHMGR tumor contains $10^8$ cells, a tumor containing 38.3 million cells would have a volume 0.38 $cm^3$, corresponding to a 7.2 mm tumor.

Engineering a 10-fold increase in limonene production by tumor cells or similar improvement in sensitivity would allow one to detect 3.8 million cells, corresponding to a tumor volume of 0.038 $cm^3$, or a 3.4 mm tumor.

Engineering a 100-fold increase in limonene production by tumor cells or similar improvement in sensitivity would allow one to detect 0.38 million cells, corresponding to a tumor volume of 0.0038 $cm^3$, or a 1.6 mm tumor.

### Supplementary Figures

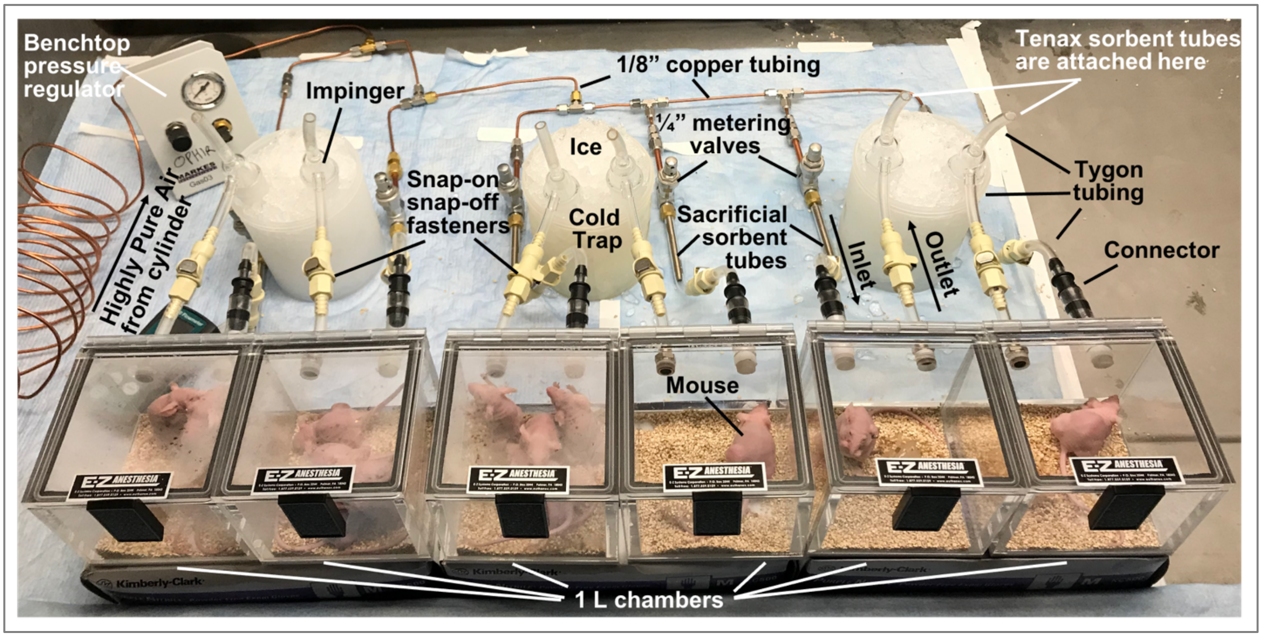


**Fig. S1. Mouse chamber/sorbent trap assembly.** Six one-liter induction chambers were operated in parallel for simultaneous mouse limonene measurements. The outlet of each chamber was connected in series via tygon tubing to a glass condenser on ice (cold trap) and then to a sorbent tube containing Tenax TA resin that traps and concentrates the VOCs. The inlet of each chamber was connected in series to a sacrificial Tenax sorbent tube, which serves to purify inflowing air, and an upstream ¼" stainless steel metering valve that individually controls air flow into each chamber. The metering valves to all six chambers were connected via reducing unions, union tees, and 1/8" copper tubing to a benchtop pressure regulator set to 5 psi, which was connected via a single copper line to a compressed gas cylinder containing highly pure air set to 20 psi. For ease of cleaning the induction chambers between experiments, the tygon connections to inlet and outlet components were interrupted by ¼" snap-on/snap-off fasteners.

| Mann-Whitney P-values | Week 1 | Week 2 | Week 3 | Week 4 | Week 5 |
|---|---|---|---|---|---|
| **HeLa-LS vs. Control** | 0.256 | 0.025 | 0.025 | 0.023 | 0.025 |
| **HeLa-LS-tHMGR vs. Control** | 0.025 | 0.025 | 0.023 | 0.023 | 0.025 |
| **HeLa-LS-tHMGR vs. HeLa-LS** | 0.049 | 0.184 | 0.105 | 0.376 | 0.049 |

**Fig. S2.** Statistical significance (Mann Whitney p-values) of limonene expression differences between HeLa-LS, HeLa-LS-tHMGR, and HeLa control mice by week. P values $< 0.05$ are highlighted in yellow.

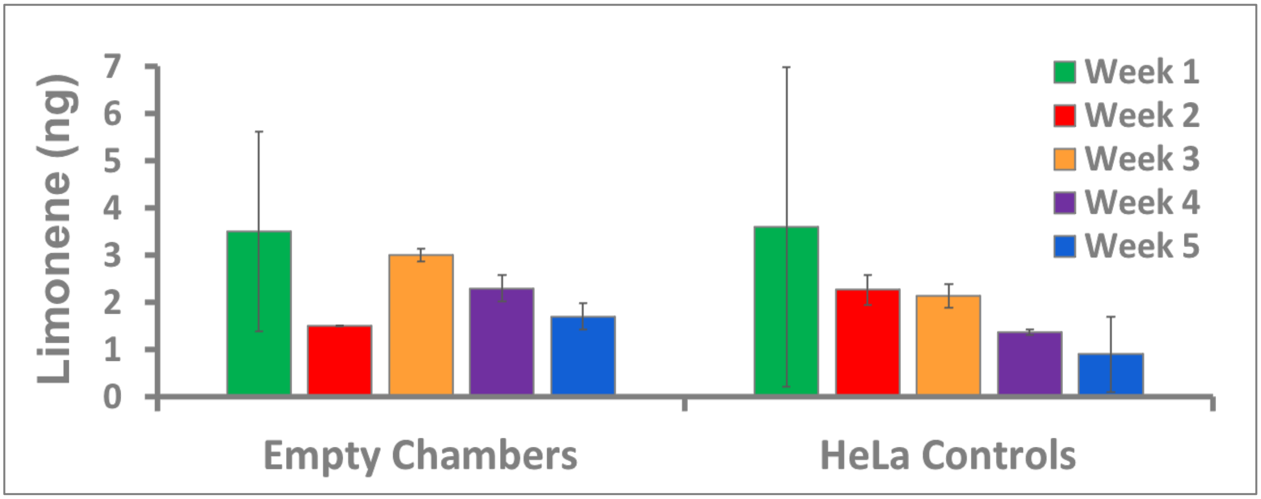


**Fig. S3. Limonene signal from empty chambers and chambers containing HeLa control mice in 10-hour sorbent trap experiments by week.** (Each bar represents mean ± SD for 3 chambers of 4 mice each; n = 12 mice).

### Supplementary Tables

| | |
|---|---|
| **Age (years)** | 30 |
| **Weight (kg)** | 70 |
| **Height (cm)** | 176.14 |
| **BMI (kg/m$^2$)** | 22.56 |
| **Hematocrit** | 0.45 |
| **Cardiac Output (mL/sec)** | 86.319 |
| **D-Limonene Clearance (L/hr)** | 1.11 |
| **2-Butanone Clearance (L/hr)** | 65.41 |

| | **Volume (mL)** | **Blood Flow (mL/sec)** | **D-Limonene Kp** | **2-Butanone Kp** |
|---|---|---|---|---|
| **Lung** | 914.414 | 86.319 | 0.90 | 0.74 |
| **Spleen** | 151.351 | 2.6578 | 4.11 | 0.78 |
| **Liver** | 1340.1 | 21.617 | 6.63 | 0.79 |
| **Gut** | - | 12.043 | 0.00 | 0.00 |
| **Adipose Tissue** | 23762.7 | 8.3458 | 16.98 | 0.26 |
| **Muscle** | 17027 | 8.9701 | 4.05 | 0.73 |
| **Heart** | 258.559 | 3.3145 | 4.47 | 0.79 |
| **Brain** | 1411.57 | 12.642 | 10.55 | 0.87 |
| **Kidneys** | 227.027 | 14.671 | 4.02 | 0.76 |
| **Skin** | 1608.11 | 3.3887 | 4.96 | 0.71 |
| **Reproductive Organs** | 26.32 | 0.0971 | 4.06 | 0.78 |
| **Bone (red marrow)** | 965.318 | 5.0855 | 11.58 | 0.58 |
| **Bone (yellow marrow)** | 2683.25 | 1.4136 | 16.98 | 0.26 |
| **Rest of Body** | 10992 | 6.7735 | 4.09 | 0.77 |
| **Hepatic Artery** | - | 6.9161 | 0.00 | 0.00 |
| **Venous Return** | 3619.61 | 86.319 | 0.00 | 0.00 |
| **Arterial Supply** | 1809.8 | 86.319 | 0.00 | 0.00 |

**Table S1. Human whole-body PBPK model parameters for an average healthy U.S. adult male.** Included in the model are organ and tissue compartments and their associated volumes and blood flow rates, as well as disposition data – including clearance and partition coefficients (*Kp*) - for limonene and 2-butanone.

| | |
|---|---|
| **Lung tissue volume (mL)** | 914.41 |
| **Lymph Volume (mL)** | 30 |
| **Mean inhalation flow (mL/s)** | 250 |
| **Fraction extracellular water volume in lung cell tissue** | 0.336 |
| **Lung tissue density (g/mL)** | 1.05 |

| | **D-Limonene** | **2-Butanone** |
|---|---|---|
| **Pulmonary Solubility at lung pH (~6.7) (mg/mL)** | 0.046 | 118.1 |
| **Vapor Diffusion Coefficient in Air (37C) (cm2/s)** | 0.1 | 0.1 |
| **Log Henry's Law Constant @ 37C (atm*m^3/mol)** | -1.55 | 0.43 |

| | **Extra-thoracic** | **Thoracic** | **Bronchiolar** | **Alveolar-Interstitial** |
|---|---|---|---|---|
| **Surface Area (sq cm)** | 450 | 290 | 2400 | 1.48E+06 |
| **Tissue Volume (mL)** | 79.53 | 56.38 | 127.3 | 510.9 |
| **Gas Volume (mL)** | 23.08 | 45.72 | 252.3 | 5675.8 |
| **Liquid (Mucus) Thickness (cm)** | 1.5E-03 | 1.1E-03 | 6.0E-04 | 6.5E-06 |
| **Epithelial Thickness (cm)** | 5.0E-03 | 5.5E-03 | 1.5E-03 | 5.81E-05 |
| **Vapor Diffusion Thickness (cm)** | 1.6 | 0.5 | 0.1 | 0.02 |
| **Liquid (Mucus) Volume (mL)** | 0.68 | 0.32 | 1.44 | 9.59 |
| **Epithelial Volume (mL)** | 2.25 | 1.60 | 3.6 | 85.70 |
| **D-Limonene Permeability (cm/s)** | 5.23E-06 | 4.76E-06 | 1.74E-05 | 4.50E-04 |
| **2-Butanone Permeability (cm/s)** | 1.48E-05 | 1.34E-05 | 4.92E-05 | 1.27E-03 |

**Table S2. Pulmonary compartment parameters.** The pulmonary compartment of the human whole-body PBPK model is divided into four sub-compartments: extra-thoracic, thoracic, bronchiolar, and alveolar-interstitial. Values for the physical properties of the pulmonary compartment as a whole (upper two tables) and of its sub-compartments (bottom table) are listed, as well as the physicochemical parameters of D-Limonene and 2-butanone in these compartments.

| Week | HeLa-LS-tHMGR | HeLa-LS |
|---|---|---|
| 1 | 49.2 | 20.6 |
| 2 | 80.6 | 44.3 |
| 3 | 134.8 | 80.5 |
| 4 | 218.0 | 129.9 |
| 5 | 332.4 | 180.8 |

**Table S3.** Calculated number of tumor cells (in millions of cells) in HeLa-LS-tHMGR and HeLa-LS mice given an estimate of $10^8$ cells/cm$^3$ of tumor tissue.

| Week | HeLa-LS-tHMGR | HeLa-LS |
|---|---|---|
| 1 | 227.7 | 89.7 |
| 2 | 372.7 | 153.0 |
| 3 | 623.3 | 328.4 |
| 4 | 1008.3 | 559.9 |
| 5 | 1537.5 | 656.5 |

**Table S4.** Predicted quantity of limonene (in ng) produced by HeLa-LS-tHMGR and HeLa-LS tumors in mice based on limonene production rates of 5.6 fg/cell/day for HeLa-LS cells and 11.1 fg/cell/day for HeLa-LS-tHMGR cells.

| Week | HeLa-LS-tHMGR | HeLa-LS |
|---|---|---|
| 1 | 7.1 | 2.6 |
| 2 | 24.8 | 16.1 |
| 3 | 28.7 | 22.3 |
| 4 | 68.7 | 57.7 |
| 5 | 92.6 | 50.3 |

**Table S5.** Measured quantity of limonene (in ng) exhaled in the breath by HeLa-LS-tHMGR and HeLa-LS mice over a ten hour period by week.

| Week | HeLa-LS-tHMGR | HeLa-LS |
|---|---|---|
| 1 | 3.1% | 2.9% |
| 2 | 6.7% | 10.5% |
| 3 | 4.6% | 6.8% |
| 4 | 6.8% | 10.3% |
| 5 | 6.0% | 7.7% |

**Table S6.** Percentage of tumor limonene that was exhaled in the breath for HeLa-LS-tHMGR and HeLa-LS mice by week.

| HeLa-LS-tHMGR | HeLa-LS |
|---|---|
| 5.2% ± 1.5% | 7.6% ± 3.1% |

**Table S7.** Percentage of tumor limonene exhaled (average over all weeks).
